# Whole genome sequences of the raspberry and strawberry pathogens *Phytophthora rubi* and *P. fragariae*

**DOI:** 10.1101/115824

**Authors:** Javier F. Tabima, Brent A. Kronmiller, Caroline M. Press, Brett M. Tyler, Inga A. Zasada, Niklaus J. Grünwald

**Affiliations:** Department of Botany and Plant Pathology, Oregon State University, Corvallis, OR, 97331, USA; Center for Genome Biology and Biocomputing (CGRB), Oregon State University, Corvallis, OR, 97331, USA; Horticultural Crops Research Laboratory, USDA-ARS, Corvallis, OR, 97330, USA

## Abstract

*Phytophthora rubi* and *P. fragariae* are two closely related oomycete plant pathogens that exhibit strong morphological and physiological similarities, but are specialized to infect different hosts of economic importance, namely raspberry and strawberry. Here, we report the draft genome sequences of these two *Phytophthora* species as a first step towards understanding the genomic processes underlying plant host adaptation in these pathogens.

## Genome announcement

More than 150 species of oomycete plant pathogens are harbored in the genus *Phytophthora* (1, 2). These organisms are highly diverse in lifestyle, host preference, and economic importance (1). Whole-genome sequences of various species in the genus *Phytophthora* have recently been published, yielding novel insights into the molecular basis of plant disease such as the identification of effector proteins that can disrupt the physiology of the host (3, 4). Here, we present the whole-genome sequences of two soil-borne *Phytophthora* sister species, *P. fragariae* and *P. rubi.* These are highly similar in morphology and physiology, but infect different hosts. Sequencing the genomes of these species will advance our understanding of the genomic mechanisms underlying host adaptation and knowledge of molecular mechanisms of plant pathogenicity (5).

Genomic DNA was extracted from *P. rubi* strain “pd0101050015038” (isolated in the Netherlands by K. Rosendhal from red raspberry) and *P. fragariae* strain CBS 209.46 (isolated in England by C.J. Hickman from strawberry). Genomes were sequenced by the Beijing Genomics Institute (BGI, Beijing, China) using the Illumina HiSeq2000 platform (Illumina, San Diego, CA) using TruSeq libraries (paired end reads, insert size of 500 bp, average read length of 90 bp). Assemblies were performed using SOAPdenovo2 (kmer size of 36) (6). Transcriptomes of both species were also sequenced by BGI using the Illumina HiSeq2000 technology (paired end reads, insert size of 500 bp, average read length of 90 bp). To obtain high quality gene calls, the transcriptomes were assembled using Trinity (7) and translated into amino-acid sequences using TransDecoder (8). To generate a reference database, gene prediction and annotation for the nuclear genomes of both species were performed using MAKER (9) trained with gene calls from other *Phytophthora* genomes including *P. ramorum, P. sojae,* and *P. infestans* (10,11). The assembled transcripts were used as RNA evidence in MAKER to improve the quality of the called gene models.

The predicted gene models were annotated using InterproScan 5 (12). The *P.fragariae* assembly was estimated to be 74 MB in size, assembled into 8,511 scaffolds with an N50 of 18,987 bp. The *P. rubi* genome was assembled into 9,434 scaffolds spanning a total length of ^~^74 Mbp and a N50 of 16,735 bp. A total of 20,448 and 23,476 gene models were found for *P. fragariae* and *P. rubi,* respectively. Both species show a high abundance of Gene Ontology terms for DNA integration, nucleic acid binding, protein binding, peptidase/helicase and hydrolase activities, integrases, endonucleases, repeats, and transporter domains.

The genomes of *P. fragariae* and *P. rubi* will provide a unique resource for these closely related soil-borne oomycete plant pathogens and the molecular mechanisms associated with plant pathogenicity. Whole-genome data have been submitted to NCBI BioProject (PRJNA375089).

## Acknowledgments

We thank Danyu Shen, Brian J. Knaus, and Jane Stewart for their support with advice and preliminary data. We also thank the Beijing Genomics Institute (BGI) for conducting the sequencing and assembly. This work was supported by funds from USDA ARS CRIS Project 2072-12220-004-00-D, 2072-22000-039-00-D, and USDA-NIFA-RAMP Project 2010-511001-21549. Mention of trade names or commercial products in this manuscript are solely for the purpose of providing specific information and do not imply recommendation or endorsement.

